# NorK, a novel norfloxacin efflux pump in *Staphylococcus aureus*

**DOI:** 10.1101/850768

**Authors:** Paul Briaud, Jessica Baude, Sylvère Bastien, Laura Camus, François Vandenesch, Karen Moreau

## Abstract

A novel efflux pump similar to Nor efflux pumps and designated as NorK was identified in *Staphylococcus aureus*. It contributes to norfloxacin resistance and presents a high level of expression in different strains. Its expression is not regulated by MgrA, unlike other genes of the *nor* gene family, suggesting that NorK could be important in the absence of expression of other *nor* genes.

---

*Staphylococcus aureus* is a major opportunistic bacterial pathogen responsible for a variety of diseases (1). Its success in causing life-threatening infections is due to its considerable production of toxins and ability to develop resistance to a wide spectrum of antibacterial compounds (2, 3). Various systems have been developed by *S. aureus* to make it resistant to these substances, such as the alteration of drug binding sites, drug inactivation and the involvement of efflux pumps.

The term multidrug resistance (MDR) pumps refers to numerous efflux pumps that are classified into two major groups: (i) ABC-type transporters, which utilize ATP hydrolysis to extrude target drugs, and (ii) secondary multidrug transporters, which exploit proton and sodium gradients as energy sources. Among this latter group, the major facilitator superfamily (MFS) has been studied extensively in staphylococcal species. In *S. aureus*, the Nor protein family belongs to the MFS and can extrude a broad range of antibiotic substances such as the fluroquinolones (FQ), norfloxacin and ciprofloxacin. Four different pumps involved in FQ resistance have been described previously (4–7). NorA and NorB are responsible for a relatively high-level of resistance to hydrophobic quinolones, whereas NorC seems to be involved in a low-level of resistance (6). The NorD substrates remain to be discovered.

Here, we report a new chromosomally encoded Nor protein called NorK, which is highly expressed independently of MgrA regulation and involved in norfloxacin resistance.

## Identification of a new putative efflux pump

In a previous study (Briaud et al., 2019), we identified 4 different *nor* related genes from RNA sequencing data, using S. *aureus* NCTC8325 as a reference genome (GenBank CP000253). Three of them have already been described: *norB* (SAOUHSC_01448), *norC* (SAOUHSC_00058) and *norD* (SAOUHSC_02762). The last gene SAOUHSC_02740 was encoded for an unknown protein and annotated as a putative drug transporter. The so-called *norK* gene is 1.401 nucleotides long (Fig. S1) and encodes for a hypothetical protein of 466 amino acids. This protein shares relatively high homology with NorB (57% identity and 76% similarity) and NorC (59% identity and 74% similarity) but low homology with the NorA and NorD pumps (16% identity and 34% similarity with NorA, 16% identity and 39% similarity with NorD). A phylogenetic analysis was conducted on Nor proteins and major chromosomically encoded MFS efflux pumps from *S. aureus* NCTC8325 using ClustalW alignment and the Maximum Likelihood method. As depicted by the cluster formed by both proteins (Fig.1), NorK is closely related to NorB and NorC. ClustalW alignment of NorK with NorB and NorC proteins revealed that highly conserved domains were shared (Fig.2). Moreover, the NorK protein presented 14 transmembrane domains predicted by the Phyre2 web portal (8), similar to those observed in the NorB and NorC pumps. Together these data suggest that NorK belongs to the same family as the Nor proteins previously described. To assess if this pump was widely distributed among *S. aureus*, BLAST analysis on a large collection of genotypically diverse bacteremia isolates (9) was performed. The results showed that 98% (123/126) harbor *norK* genes (identity and coverage >90%).

**FIG. 1.**
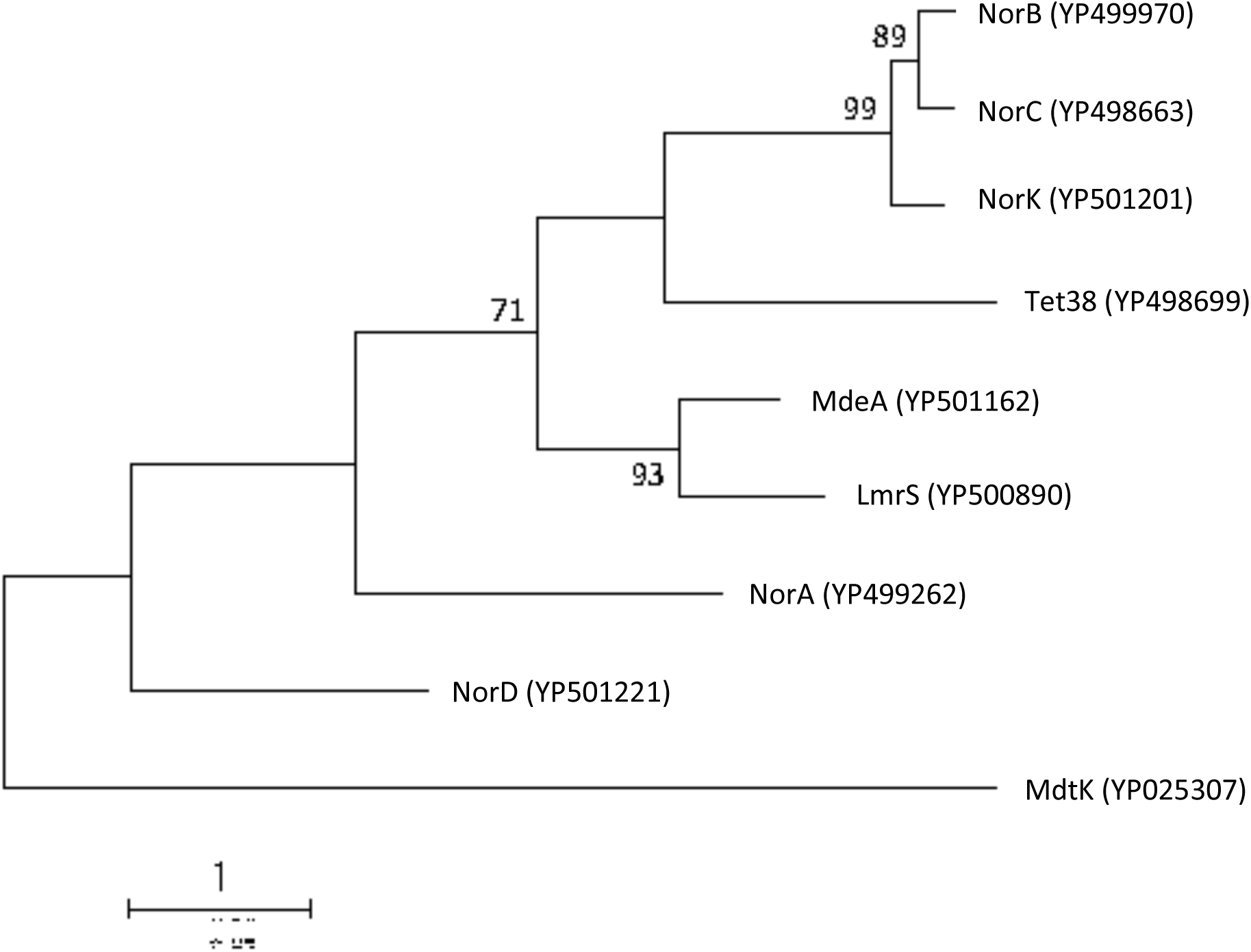
Phylogenetic tree of *S. aureus* chromosomally encoded major MFS efflux pumps. Accession numbers from the *Staphylococcus aureus* NCTC 8325 strain are indicated in parentheses. Proteic sequences were aligned by the ClustalW method and the tree was built using the Maximum Likelihood method based on the Le Gascuel 2008 model (14) with a bootstrap of 10000. The tree with the highest log likelihood (−7115.58) is shown. The percentage of trees (>70) in which the associated taxa clustered together is shown next to the branches. The Multi-Antimicrobial Extrusion Protein (MATE) MdtK sequences from *E. coli* (KAB2830459) was used to root the tree.

**FIG. 2.**
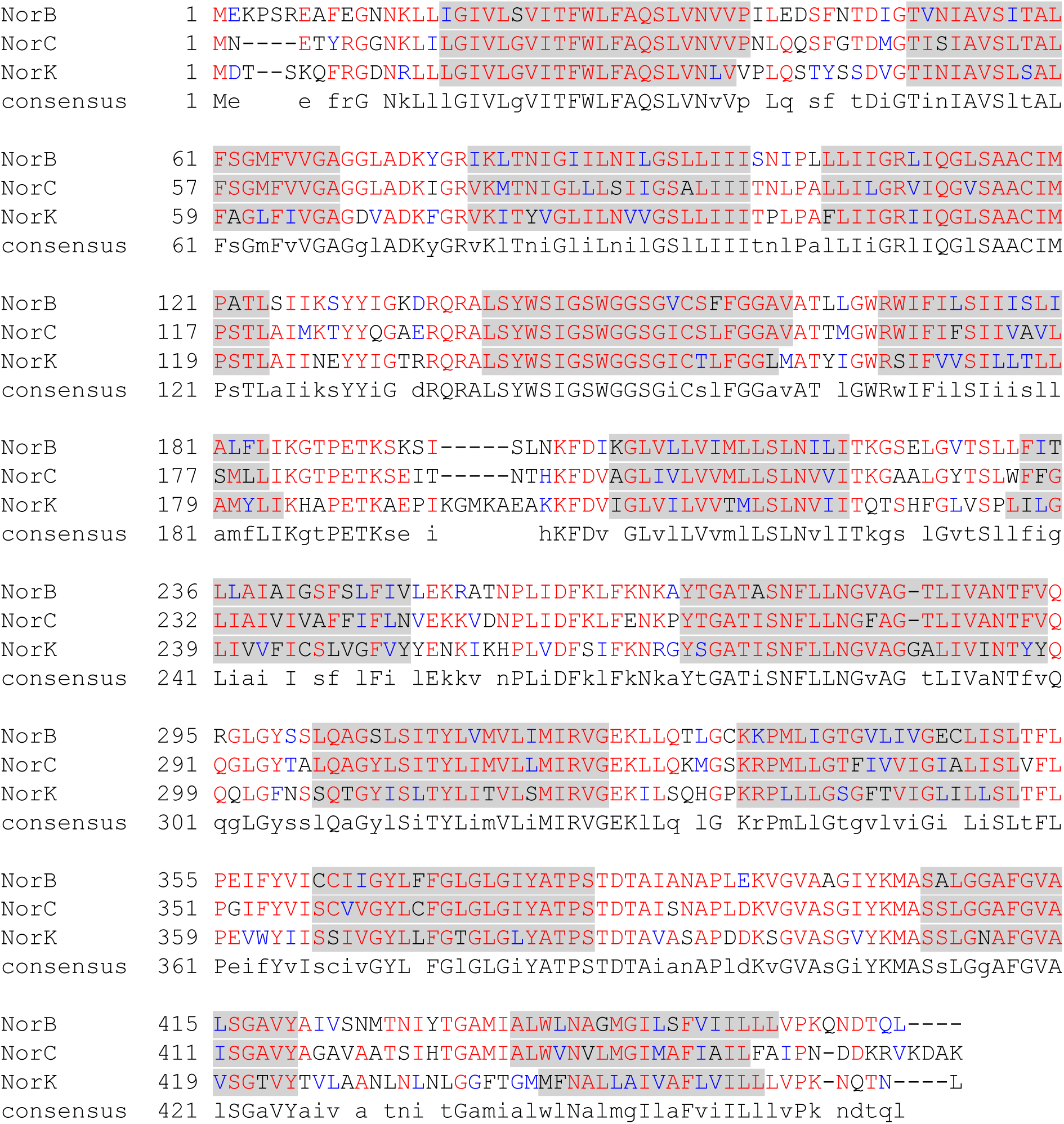
Amino acid sequence alignments of *Staphylococcus aureus* NCTC 8325 NorB (YP 499970), NorC (YP 498663) and NorK (YP 501210) proteins. Alignment was done using ClustalW. Identical amino acids are shown in red, similar amino acids in blue. Grey boxes represent predicted transmembrane domains by Phyre2 web portal (8).

The expression of *norK* genes was explored by RT-qPCR (protocol in supplementary data) on 5 different laboratory strains of *S. aureus* and compared to *norA, B, C* and *D* gene expressions. The *norK* transcript was consistently detected in all the strains tested at a level similar to *norA* and *norD* genes (Fig. 3 and S2), supporting functionality.

**FIG. 3.**
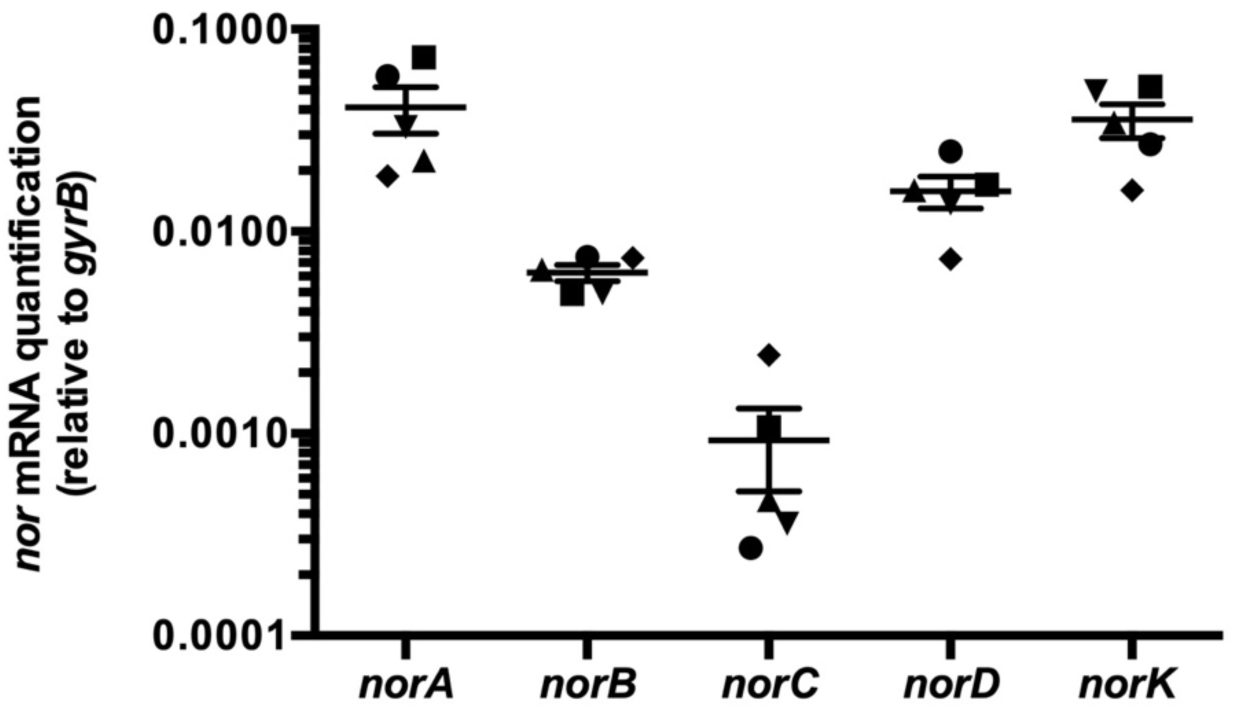
*nor* genes expression. *S. aureus* strains RN6390• (14), HG001▾ (15), SH1000▪ (16), Newman♦ (16), SF8300▴ (17), were cultivated in BHI for 8 hours at 37°C with shaking (200rpm). At 4 hours of culture, RNAs were extracted and *nor* gene expression was monitored by RT-qPCR. The results are represented as the mean +/- standard error. Quantification after 2, 6 and 8 hours of culture are presented in figure S2

MgrA can act as a repressor or an activator of *norA, norB and norC* genes (5–7) depending on the RsbU strain background and as a result of its phosphorylation state (22). We thus explored *norK* gene expression in the Newman wild-type (WT, rsbU+) and the Newman Δ*mgrA* strain (11). *mgrA* deletion had no impact on *norK* expression while increased *norA* and *norC* and decreased *norB* expression were observed as expected for a RsbU positive background (Fig. 4) (10, 12).

**FIG. 4.**
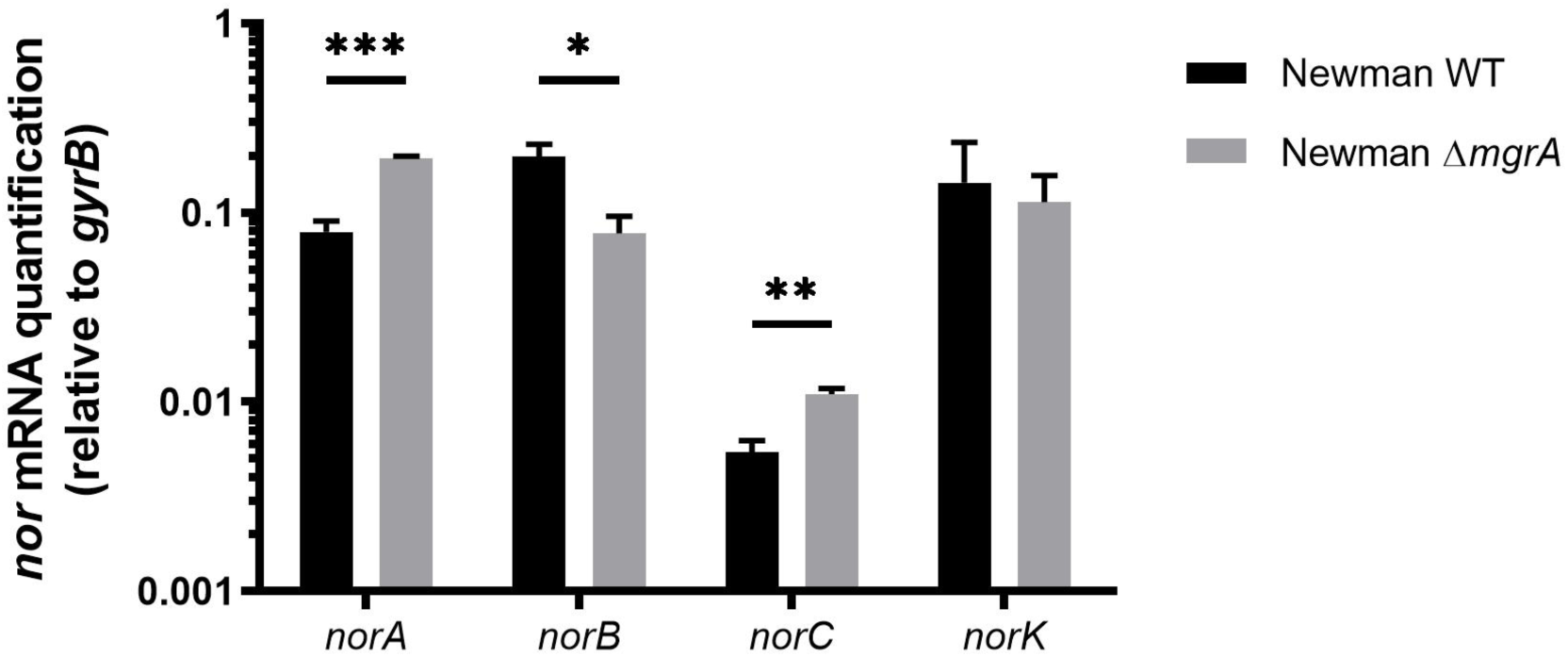
MgrA regulation of *nor* gene expression. Newman wild-type (black) and Newman Δ*mgrA* (grey) strains were cultivated in BHI at 37°C for 8 hours. At 4 hours of culture, RNAs were extracted and *nor* gene expression was monitored by RT-qPCR. The results are depicted as mean + standard error from three independent experiments. Statistical analyses were performed by using an unpaired t-test (* P<0,05, ** P<0,001, *** P<0,0001).

## NorK efflux activity and involvement in antibiotic resistance

To assess the functionality of NorK, thus its efflux activity, an overexpressing *S. aureus* strain was built (protocol in supplementary data) and an ethidium bromide efflux assay was performed as previously described (13). The rate of fluorescent loss in *norK*-overexpressing strains was significantly higher compared to the WT strain (Fig. 5). Antibiotic efflux was tested by time-kill assay in the presence of a representative anti-staphylococcal fluoroquinolone, norfloxacin, as previously described (14). Wild-type and *norK*-overexpressing strains were cultivated for 8 hours in Mueller-Hinton broth supplemented with norfloxacin at ½MIC (0.19µg/mL), MIC (0.38µg/mL) and 2MIC (0.76µg/mL). At 2, 4, 6 and 8 hours, bacteria were enumerated by plating on TSA. The survival rates of the *norK*-overexpressing strain were significantly higher (between 0.5 log and 1 log CFU/ml) than those for the WT strain.

**FIG. 5.**
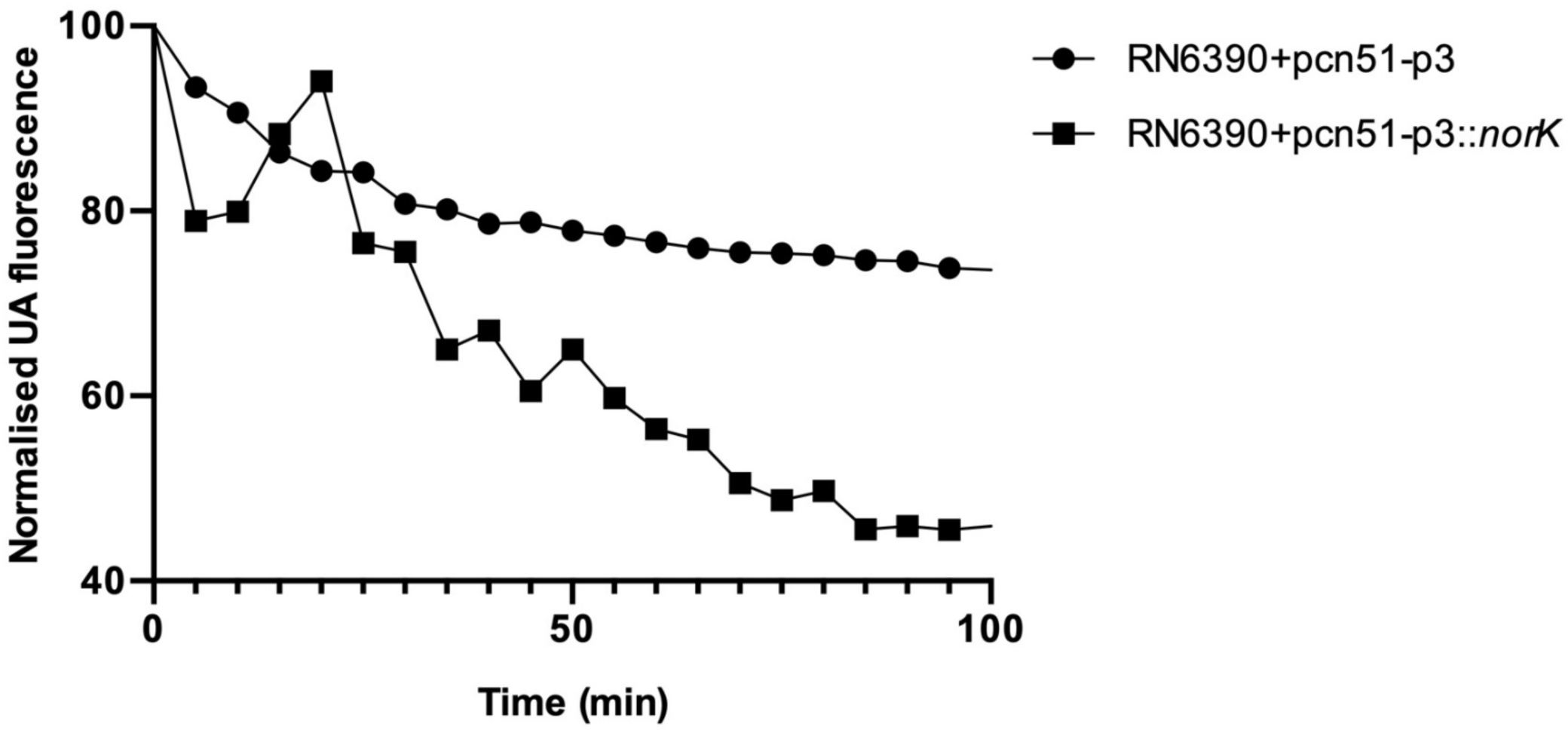
Quantification of ethidium bromide efflux over time by *S. aureus* RN6390 WT (•) and *norK* overexpressing (▪) strains. Bacteria were resuspended in efflux buffer with EtBr (2µg/mL) and reserpine (25µg/mL), to prevent efflux. After 30min, bacteria were washed to remove reserpine and EtBr efflux was monitored by measuring the decrease of fluorescence. One experiment representative of three independent experiments is depicted.

**FIG. 6.**
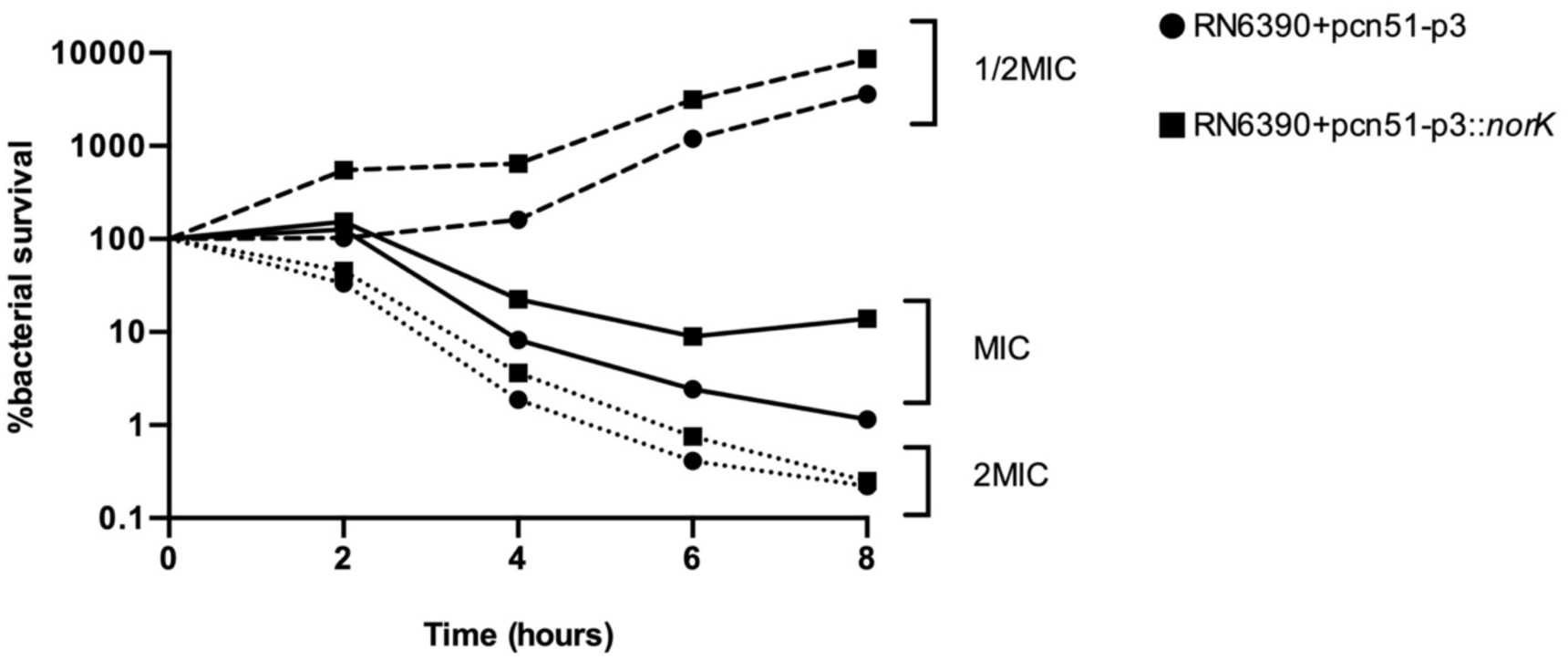
Time–kill curves of *S. aureus* RN6390 WT (•) and *norK* overexpressing (▪) strains. Bacteria were grown in Mueller-Hinton broth (MHB) for 8 hours. Bacteria were resuspended at 1.10^6^ CFU/mL in MHB supplemented with norfloxacin at ½ MIC (0.19µg/mL), MIC (0.38µg/mL) and 2MIC (0.76µg/mL). Bacteria were plated on MHA every two hours to count living cells. The results are indicated as the percentage of survival rate calculated by dividing the number of bacteria at a time point by the number of bacteria at T0. One experiment representative of three independent experiments is depicted.

In conclusion, NorK is a new member of the *nor* gene family and contributes to *S. aureus* norfloxacin resistance. NorK is widely distributed within *S. aureus* strains and its high level of expression may point to an important role in antibiotic resistance and fitness during the colonization of infectious niches. Moreover, the MgrA-independent regulation of *norK* indicates that its expression could be induced even though other *nor*-regulated genes were not expressed. Further studies will be necessary to explore the expression profiles of *nor* genes in different conditions (e.g., antibiotic stresses, infectious niches).

## Acknowledgments

This work was funded by Inserm. P. Briaud was funded by the French Ministry of Education and Research, and L. Camus was funded by the Fondation pour la Recherche Médicale (grant number ECO20170637499).

## Authors’ contributions

PB, JB, LC, FV and KM contributed to the design of the study. PB and JB conducted the experiments. SB conducted and analyzed all the bioinformatics works. PB and KM collected the data and wrote the first draft of the manuscript. All the authors contributed to manuscript revision and approved the submitted version.

None of the authors have any conflict of interest to report.

